# Feel the Beat: The Impact of Rhythms on Dynamic Visual Search

**DOI:** 10.64898/2025.12.20.695695

**Authors:** Talya Shlesinger-Arad, Flor Kusnir, Leah Snapiri, Nir Shalev, Ayelet N. Landau

## Abstract

Rhythms are everywhere – we encounter them daily within our environment and produce them regularly through our movements, speech and sensory systems. In both cognitive and clinical contexts, external rhythms have been shown to enhance performance when they are aligned with, predict, or entrain stimuli onsets or relevant moments in time. In this study, we aimed to understand the impact of external and internal rhythms on performance in a task devoid of rhythmic temporal structure. We thus investigated the differential impact of auditory beats (1-, 2-, 5-Hz, or no beat) on detection in a non-rhythmic, dynamic search task requiring the continuous deployment of attention. We additionally assessed whether the influence of auditory beats on detection was modulated by individuals’ rhythmic preferences as measured by their spontaneous motor tempo. Overall, we found that auditory rhythms enhanced visual task performance, even in the absence of rhythmic structure within the measured task, improving both detection accuracy and response speed. This enhancement was tempo-dependent, with the strongest facilitation observed at 2 Hz. The magnitude of the observed benefits varied with individuals’ spontaneous motor tempo, such that participants with slower SMTs exhibited the greatest performance benefits due to the presence of auditory rhythms. We propose that rhythmic stimulation may serve as a powerful scaffold for sustained performance in the context of dynamic and temporally unpredictable environments.

## INTRODUCTION

Rhythms are embedded in the dynamic stimuli we encounter daily, including biological motion (Fraisse & Deutsch, 1982; Lakatos et al., 2019; Snapiri et al., 2025), speech, and music (Inbar et al., 2020, 2025; Poeppel & Assaneo, 2020). The temporal structure of the environment shapes our ability to anticipate sensory input, modulating perception and helping us temporally coordinate our responses (Haegens & Zion Golumbic, 2018; Herbst & Landau, 2016; Jones, 2010; Large & Jones, 1999; Nobre & Van Ede, 2018; Shalev et al., 2019). In addition to external rhythms in the environment, rhythms are also spontaneously generated by the brain and the body. For example, spontaneous motor tempo (SMT) – which are generated by the motor system – is a consistent trait within individuals, stable across both time and a variety of spontaneous motor tasks (e.g., tapping, walking, bouncing) (Engler et al., 2024; Rimoldi, 1951; Snapiri et al., 2025). In contrast, *across* individuals as well as across the lifespan, there is substantial variability in the rhythm of these naturally paced behaviours (Desbernats et al., 2023; McAuley et al., 2006). Recently, this stable, individual trait – SMT – has been shown to account for individual differences in perception, memory, and behaviour (Hine et al., 2025; Roman et al., 2023; Snapiri et al., 2023; Zamm et al., 2015, 2016, 2018), suggesting that SMT affects cognition beyond merely the motor system.

Due to its ability to prime the motor system and provide temporal stability, rhythm has increasingly been used as a therapeutic tool in motor and speech rehabilitation (Fujii & Wan, 2014; Nombela et al., 2013; Thaut et al., 2015; Thaut & Abiru, 2010). Auditory-motor entrainment (achieved, for example, by using a metronome as individuals engage in movement) is supported by physiological and functional connections between the auditory and motor systems and has been successfully applied to a number of clinical populations, e.g., Parkinson’s, stroke, traumatic brain injury, cerebral palsy, and hemiparetic patients (Bégel et al., 2018; Braun Janzen et al., 2022; Daigmorte et al., 2022; Puyjarinet et al., 2017). For example, Rhythmic Auditory Stimulation (RAS) is widely used in Parkinson’s disease and stroke to improve gait in patients undergoing gait training. RAS is implemented through having patients synchronize their steps to externally-presented, rhythmic auditory cues that function as an external timekeeper, entrain motor output, and compensate for impaired internal timing mechanisms (McIntosh et al., 1997; Thaut et al., 1996, 1997; Yoo & Kim, 2016). Additionally, Patterned Sensory Enhancement (PSE), another treatment that uses the rhythmic elements of music to provide temporal, spatial, and dynamic cues for movements, has also shown benefits in reducing movement variability and normalizing the velocity profiles of the paretic arm after stroke (Bharathi et al., 2019; Braun Janzen et al., 2022; Thaut & Abiru, 2010). Similarly, rhythmic interventions are used in developmental disorders and rehabilitation. In dyslexia, regular metrical primes are used to facilitate syntax and phonological processing via neural entrainment (Schön & Tillmann, 2015). In Attention Deficit Hyperactivity Disorder, rhythmic auditory stimulation and background music are employed to improve inhibition control and optimize arousal levels for academic tasks (Abikoff et al., 1996; Park et al., 2013). Furthermore, in aphasic patients, pulse-based rhythmic interventions such as Melodic Intonation Therapy utilize hand-tapping to facilitate speech motor output (Fujii & Wan, 2014; Lim et al., 2013; Wiltshire et al., 2024).

In cognitive research, the effects of rhythm have been explored primarily through two complementary experimental approaches: rhythmic facilitation and background music. In rhythmic facilitation paradigms, rhythms are presented auditorily or visually, and goal-relevant targets appear either on-beat or off-beat. This design allows researchers to determine whether temporal alignment between rhythmic structure and target onset confers behavioral advantages (Bauer et al., 2015, 2021; Cravo et al., 2013; Shalev & Nobre, 2022; Snapiri et al., 2023). Thus, this literature utilises a rhythmic context that is highly task-relevant. In contrast, studies examining the effects of background music on cognition have focused on the impact of tempo orthogonal to the measured task (e.g., in attention, problem solving, memory; see Cheah et al. (2022) for a review), with several studies suggesting an impact (Angwin et al., 2018; Kiss & Linnell, 2021; Mendes et al., 2021; Wu & Shih, 2021). For example, Husain et al. (2002) reported enhanced accuracy on a spatial reasoning task, following exposure to fast-tempo music compared to slow-tempo music (e.g., identifying unfolded paper shapes). In contrast, Thompson et al. (2012) found that fast and loud background music hinders reading comprehension. A more recent study supports a link between arousal (induced by background music) and attentional state (i.e., decreases in mind wandering and increases in task focus) (Kiss et al., 2024). Nonetheless, the findings in this line of research are inconclusive, reporting both beneficial and detrimental effects on performance without a clear understanding of which parameters influence this discrepancy.

One way to reconcile the apparent conflict among findings is to consider individual differences in response to rhythmic stimulation. One such source of interindividual variability could stem from individual preferred event rates. For example, sensorimotor synchronization is reported to be less accurate and less stable in slow tempi or in cases where the tempo in use is slower than an individual’s SMT (Varlet et al., 2012). In addition, several studies indicate that individuals achieve optimal performance when operating at tempi that align with their SMT (McAuley et al., 2006; Zamm et al., 2015, 2016, 2018). In line with these observations, rehabilitation studies sometimes measure an individual’s SMT prior to the intervention in order to determine a personalised tempo for learning or training (Bella et al., 2017; Benoit et al., 2014; Cochen De Cock et al., 2021; Leow et al., 2014). It is therefore possible that, in non-rhythmic tasks, the presence of background rhythms may exert differential effects across individuals depending not only on the specific modulation tempo but also on each person’s rhythmic preferences.

In order to better understand the impact of external and internal rhythms on performance, our investigation focuses on the differential impact of beats on a non-rhythmic, dynamic task that requires continuous deployment of attention. We address two main questions: (i) what are the effects of an external tempo on attention and performance in a non-rhythmic task? Importantly, and as is most often the case in our daily life, the external tempo here was unrelated to the task, conferring no information about the onset of upcoming targets. (ii) We also asked how these effects are modulated by an individuals’ rhythmic preferences, as measured by SMT. To this end, we employed a dynamic visual search task requiring participants to monitor a set of stimuli continuously fading in and out of the display. Participants were instructed to detect targets by tapping on them before they disappeared from the screen. Unbeknownst to them, half of targets appeared at predictable onsets and approximate locations across trials, and the other half appeared randomly (in both time and space) (Boettcher et al., 2022; Shalev & Nobre, 2022). The experimental design was selected for its continuous nature. This task was combined with external, task-irrelevant auditory stimulation in order to examine the primary aim of the study. The dynamic search task permits the extraction of multiple indices relevant to the influence of external rhythms, including general performance measures (accuracy and response speed) as well as task-specific indicators of implicit learning of temporal regularities.

While performing the dynamic search task, participants listened to ambient background noise (Brownian noise spectrum) that contained various environmental sounds (birds and other nature sounds). Critically, within this pleasant and familiar soundscape, we embedded a beat with varying frequencies (1 Hz, 2 Hz, 5 Hz, or no beat). Through this setup, we measured how auditory rhythms unrelated to the task at hand impact overall detection performance. Additionally, we explored whether the impact of external rhythms is dependent on individuals’ internal rhythms. Prior to the dynamic search task, we thus measured individuals’ spontaneous motor tempo using a simple tapping task and tested whether individual differences in SMT further modulate the impact that rhythms have on performance.

## METHODS

### Participants

Seventy-one participants (54 males, mean age = 23.1 ± 2.4, 12 left-handed) recruited from the Hebrew University of Jerusalem participated in the experiment. The sample size was determined based on previous individual differences studies (Siegelman et al., 2017; Snapiri et al., 2023). All participants had normal or corrected-to-normal vision and no reported history of major medical illness, head trauma, neurological or psychiatric disorders. We recruited participants using the university system for participant recruitment. We obtained written informed consent from all participants before the experimental session. Monetary compensation or course credit was provided for participation in the study. All experimental procedures were approved by the Ethics Committee of the Hebrew University of Jerusalem.

### Experimental Tasks

Throughout the following experimental tasks, participants were seated comfortably on a sofa in a quiet, well-lit room. The room was designed to mimic a leisurely space, similar to a living room, in order to create an “everyday” atmosphere for the experiment.

### Spontaneous Motor Tempo Task

In order to measure participants’ spontaneous rhythmic preferences, we first administered the spontaneous motor tempo task (Fraisse & Deutsch, 1982; McAuley et al., 2006) by instructing individuals to tap on an Android phone screen, with the index finger of their dominant hand, for 60 seconds at a consistent and comfortable tempo. We recorded tapping times using a touch-sensitive app developed in-house. We placed the phone on a clipboard positioned on the participant’s knees.

### Dynamic Search Task

The task is depicted in Figure 1 and was performed on a 10″ Microsoft Surface tablet (28 × 17 cm; 60 Hz refresh rate) running Linux. Participants placed the tablet on their lap at a comfortable position and distance and wore headphones. The experimental script was generated using Psychopy and displayed on the tablet via the Pavlovia.org online experimental platform (Peirce et al., 2019). The experimental design was adapted and modified from a previously used experiment (Boettcher et al., 2022).

**Figure 1.**
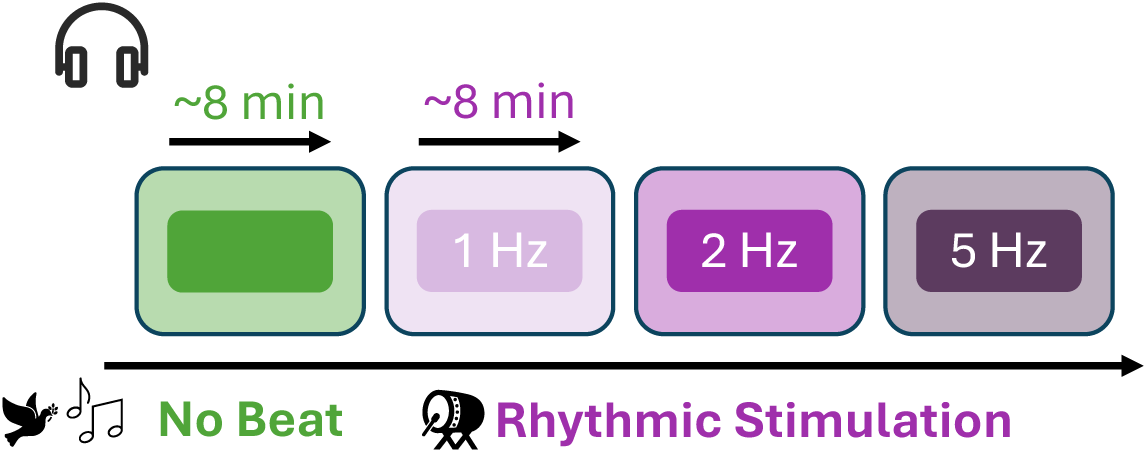
Experimental design. Participants engaged in four separate blocks. While participants performed a visual dynamic search task, they wore headphones and heard environmental and ambient background sounds. Blocks differed by their rhythmic stimulation condition and could either have no beat (depicted in green; always presented either first or last) or a beating drum with a 1 Hz, 2 Hz or 5 Hz beat frequency (depicted in shades of purple). The order of the rhythmic stimulation blocks was counterbalanced across participants.

Participants were instructed to find and tap on eight targets in each trial, which lasted approximately 14 seconds. Each trial comprised eight targets and sixteen distracting stimuli. Stimuli were non-grapheme symbols, of which targets were defined by a dark blue color and distractors by a light blue color. Any symbol could be a target or distractor on any given trial, i.e., targets and distractors were purely defined by color.

Stimuli slowly faded in and out of the search display independently over the entire trial duration. They faded in over 1 second, stayed visible for another 0.5 seconds, and then faded out over 1 second. The search display background consisted of a 1/F static noise patch generated for each trial.

Each stimulus could appear in any of the four display quadrants as long as it did not overlap with another stimulus. Participants were required to tap on the location of the targets upon detection. Tapping did not alter the appearance of the stimulus, and no immediate feedback was given. Participants were instructed to tap with their finger on the targets without a specific limit on the number of taps per target or trial.

### Target Predictability

The dynamic visual search task manipulated the spatiotemporal predictability of target stimuli (Figure 2a and 2b). Among the eight target stimuli, four were predictable, appearing with the same temporal onset and within the same quadrant on every trial. However, their specific location within the quadrant varied, as did the specific symbol used. Onset times for predictable targets were pre-assigned by dividing the time period from 0.6-10 seconds after trial onset into four equally sized time windows (“time bins”). Each time bin was further subdivided into six equally spaced onset times (twenty-four bins in total). Within each bin, one predictable target was assigned to one of the six onsets (no more than one predictable target for each bin). These interval-quadrant associations remained fixed for each participant throughout each block, i.e., they were reset at the beginning of each block. The remaining four random targets and sixteen distractors were then assigned to the remaining onset times randomly in each trial, with an additional random time jitter ranging from −0.5 to 0.5 seconds. This arrangement ensured an even distribution of targets throughout the trial and prevented an excessive number of target events at any given time. The combination of random targets among predictable targets, as well as the spatial uncertainty of random targets, prevented the learning of regularities based solely on sequences or spatial locations.

**Figure 2.**
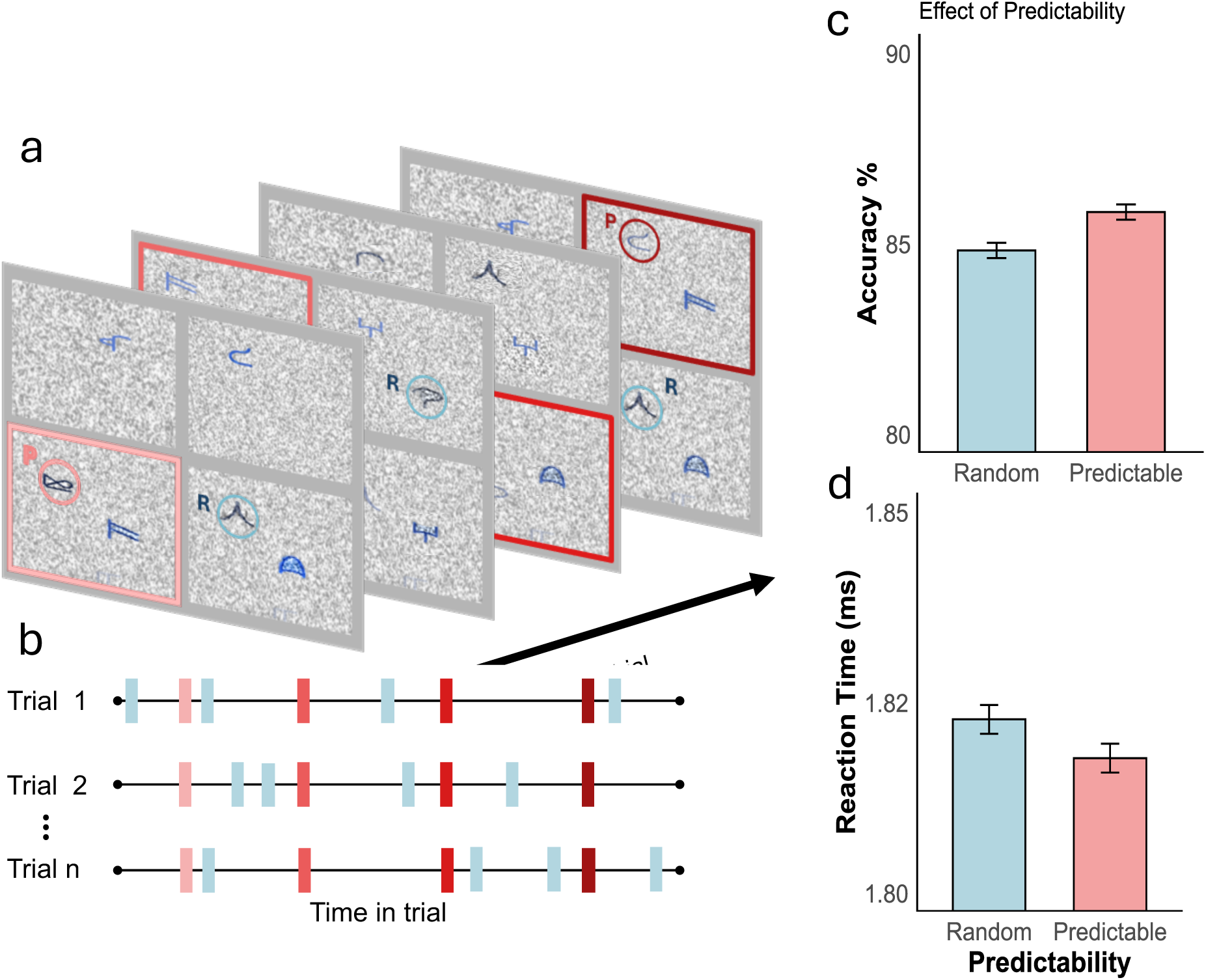
Trial Schematic. (a) In each trial, participants searched for eight dark blue symbols among light blue distractors. For visualization purposes, a pink circle indicates the first predictable target, and a turquoise circle indicates the first random target in the trial. Darker red hues indicate the quadrant and position in which the three subsequent predictable targets appeared. Targets and distractors appeared and disappeared over the course of the entire trial. (b) The time course of a trial is depicted with the onset of trial events represented as rectangles (pink scaled rectangles represent predictable targets while turquoise rectangles represent unpredictable targets). Predictable targets always appeared at the same time and within the same quadrant (although the exact location within the quadrant varied). Unpredictable targets had variable onsets and could appear anywhere. (c) Effect of target predictability on detection performance in our sample. Detection performance for predictable targets was significantly higher than for random targets. (d) This is reflected in the RTs, which were faster for predictable targets compared to random ones. Error bars on plots c and d represent within-subject standard error (Cousineau-Morey method) (Morey, 2008), using the “summarySEwithin” function in R from the Rmisc package (Hope, 2013).

### Distractors

Distractors were symbols that were identical to the targets in every aspect except for color. Distractors were presented in light blue, while targets were shown in dark blue. Importantly, the distractors had no temporal or spatial predictability.

### Experimental Procedure

Participants underwent ten practice trials to acquaint themselves with the task before completing four blocks comprising 35 trials each. Each trial contained eight targets (4 predictable and 4 random), totaling 280 targets (140 predictable and 140 random) per participant, excluding practice trials. In addition, each trial contained 16 distractors that participants were instructed to ignore, though they were not penalized for tapping on them. After each 14-sec trial, a screen was displayed with feedback indicating how many targets (out of the eight) participants had detected. The feedback remained on the screen until the participant tapped to proceed, allowing them to control the pace of the experiment. The experimental block lasted approximately 8 minutes. At the end of a block, an experimenter started the next one, encouraging a short break between blocks.

### Auditory Rhythmic Stimulation

During each block and while performing the dynamic search task, participants wore headphones and listened to auditory drum-like beats, embedded in an ongoing, non-rhythmic background sound consisting of brown noise combined with bird and rain sounds. These stimuli were designed in-house using Audacity. Beats were constructed based on three frequencies: 1, 2, and 5 Hz. These frequencies were chosen based on previous SMT research (Desbernats et al., 2023; Hammerschmidt et al., 2021; Moelants et al., 2002) so as to represent slow, medium, and fast tempos to listeners. These studies showed that the population’s tempo of tapping is centered around 2 Hz, with 1 Hz and 5 Hz serving as the extreme values of the typical tapping range. During each experimental block, participants heard one of the three beats (1, 2, 5 Hz) or only the background sounds without an embedded beat (referred to as “no-beat”). The no-beat sounds were always presented either in the first or last block (counterbalanced across participants), and the other three beat-blocks were randomly shuffled for each participant.

### Data Analysis

To examine participants’ ability to detect targets as a function of external rhythmic stimulation, individual spontaneous tempo (SMT) and predictability of target onsets, we performed a mixed effect logistic regression (Jaeger, 2008). In separate analyses, we examined two dependent measures: detection accuracy and reaction times. Correct responses were coded as 1 and incorrect responses as 0. Our independent measures included ‘rhythmic stimulation’, ‘SMT’, ‘predictability’ and all the interactions between them as fixed effects. We also included random intercepts for participants. The two categorical predictors, rhythmic stimulation (no beat, 1, 2 and 5 Hz) and predictability (random, predictable) were ‘treatment coded’ (reference levels: ‘no beat’ and ‘random’). The continuous predictor, SMT, was mean-centred.

To assess the contribution of a specific predictor on the model, we performed a likelihood ratio test between the full model and a nested model that excluded that specific predictor. We repeated this for each predictor separately. In what follows, we report the Bayesian Information Criteria (BIC) difference, chi-square values and significance level for each of our three predictors (Meteyard & Davies, 2020).

To obtain comparisons of interest that were not covered by the contrast structure included in the model, we utilized the ‘emmeans’ package in R (Lenth & Piaskowski, 2025). With this package, we calculated the log odds difference between rhythmic stimulation conditions (beats vs. no beat) as well as target predictability (predictable vs. random). Then, we tested these differences for significance using the Wald Z-test and transformed the log odds difference into odds ratio. For each comparison, we report odds ratio (OR) with confidence intervals and adjusted significance levels using false discovery rate correction for multiple comparisons (Benjamini & Yekutieli, 2001).

### Data Preprocessing

We excluded participants with low accuracy or high false alarms by computing the mean of each metric across participants and identifying outliers using the interquartile range method (IQR). Two participants were excluded due to low accuracy (59% and 61%), and five participants were excluded due to high mean false alarms (9-13 false alarms per trial). False alarms were defined as instances in which participants touched one of the distractors. An additional participant was excluded due to missing SMT data. Our final sample included 59 participants.

## RESULTS

### Performance in visual search task is dependent on rhythmic stimulation

We first examined whether rhythmic stimulation impacted participants’ ability to detect targets. We performed a model comparison between the full model and a nested model which excluded the main effect of rhythmic stimulation. We found that rhythmic stimulation contributed significantly to the full model, meaning that relative to no beat, an external rhythmic beat impacted detection accuracy in the dynamic visual search task (ΔBIC = 70, χ^2^ (12) = 63.12, *p* =<.001; 85.67±6.98% vs. 84.25±8%).

To further characterize how performance varied by beat (compared to no beat) as well as between beat rhythms, we obtained log ratios between the different tempi (1, 2, 5 Hz, and no beat). We found that, compared to the odds of detecting a target without a beat, the odds of detecting a target with the 1 Hz beat were 1.09 more likely (*OR* = 1.09, 95% CI: [1.03 1.16], *p* = 0.0086; 85.38±7.7% vs. 84.25±8%); the odds of detecting a target with the 2 Hz beat were 1.19 higher (*OR* = 1.19, 95% CI: [1.12 1.27], p < 0.0001; 86.42±6.91% vs. 84.25±8%); and the odds of detecting a target with the 5 Hz beat were 1.07 higher (OR = 1.07, 95% CI: [1.01 1.14], p = 0.025; 85.21±6.34% vs. 84.25±8%). We additionally observed improved detection performance with the 2 Hz beat compared to the 1 and 5 Hz stimulation (1 Hz vs. 2 Hz: *OR* = 1.09, 95% CI: [1.03 1.16], p = 0.0086, 85.38±7.7% vs. 86.42±6.91%; 5 Hz vs. 2 Hz: OR = 1.11, 95% CI: [1.04 1.18], p = 0.0034, 85.21±6.34% vs. 86.42±6.91%). In summary, our results show a rhythmic modulation on detection performance, such that all rhythmic conditions improved performance relative to having no rhythm; and, additionally, the 2 Hz beat benefited performance relative to the other rhythmic stimulation beats (1 and 5 Hz).

We observed a complementary pattern in the analysis of RTs. We found that rhythmic stimulation contributed significantly to the full model, meaning that relative to no beat, an external rhythmic beat impacted RTs (ΔBIC = 210, χ^2^(12) = 341.19, *p* =<.001; 1.82±0.2% vs. 1.84±0.17%). We again further characterized how RTs varied by beat (compared to no beat) as well as between beat rhythms. To this end, we obtained estimates of the contrasts between all pairs. We found that, compared to the no beat, the 1 Hz and 2 Hz conditions resulted in significantly faster RTs (1 Hz vs. no-beat: estimate: −0.011, [-0.014 −0.0088], *p* < .0001; 1.81±0.21% vs. 1.84±0.17%; 2 Hz vs. no-beat: estimate: 0.016, [-0.019 −0.014], *p* <.0001; 1.79±0.2% vs. 1.84±0.17%), while the 5 Hz rhythm hindered RTs (estimate = 0.0028, [0.00012 0.0054], *p* <0.05; 1.85±0.18% vs. 1.84±0.17%]. Between rhythms (1, 2, 5Hz), the 2 Hz again benefitted performance most relative to the other beats (1 vs. 2 Hz: estimate = −0.0055, [-0.0082 −0.0029], *p* < 0.001, 1.81±0.21% vs. 1.79±0.2%; 5 vs. 2Hz: estimate = −0.20, [-0.022 −0.017], *p* <.0001, 1.85±0.18% vs. 1.79±0.2%; 5 vs 1Hz: estimate = −0.014, [-0.022 −0.017], *p* <.0001, 1.85±0.18% vs. 1.81±0.21%).

**Figure 3.**
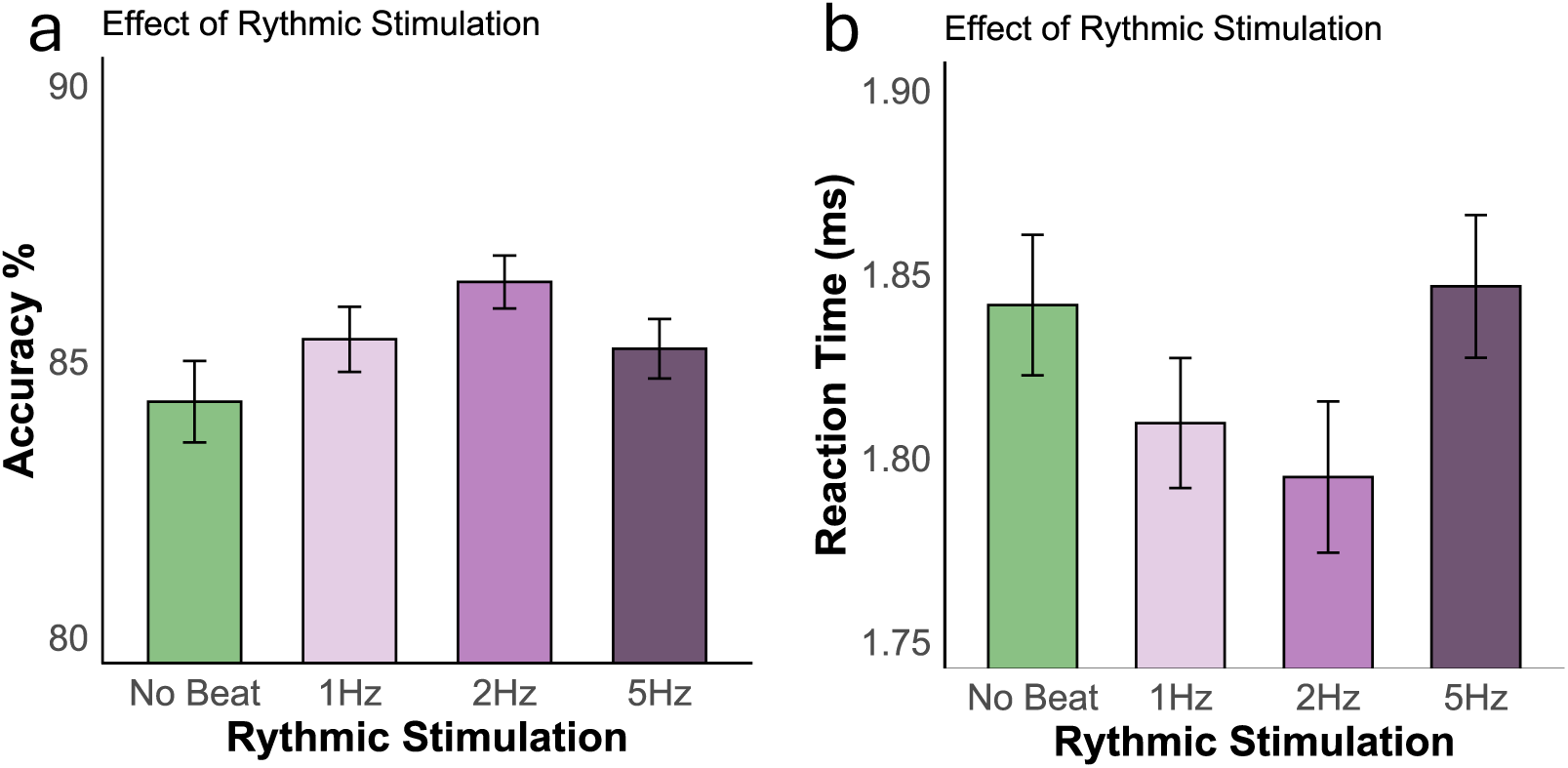
Impact of rhythmic stimulation on detection performance and response speed. (a) Compared to the no-beat condition, detection accuracy was significantly better when participants heard background rhythms. The medium tempo (2 Hz) conferred the greatest benefits to detection accuracy compared to the slow (1 Hz) and fast (5 Hz) tempi. (b) A complementary pattern was observed in the RTs of the same participants. Error bars on the plots represent within-subject standard error (Cousineau-Morey method) (Morey, 2008), using the “summarySEwithin” function in R from the Rmisc package (Hope, 2013).

### Individuals’ spontaneous motor tempo is linked to the impact of rhythmic stimulation

To assess whether the impact of external rhythms on detection performance is influenced by individuals’ spontaneous tempo, we compared the full model and a nested model which excluded the interaction between external rhythmic stimulation and personal spontaneous tempo. We found that the impact of external rhythms on performance was indeed linked to personal tempo (ΔBIC =39, χ^2^ (6) = 27.027, *p* =<.001).

We further examined this interaction by first estimating, for each external rhythm separately, how detection performance varies as a function of spontaneous tempo (see Figure 4, below). To this end, we estimated the slopes of the linear trend between detection performance and spontaneous tempo for each external rhythm separately (using the emtrends() function of the emmeans package). Then we statistically compared these slopes in order to assess whether detection performance is modulated by SMT equivalently across external rhythms, or whether some external rhythms influence the relationship of personal tempo on detection performance.

**Figure 4.**
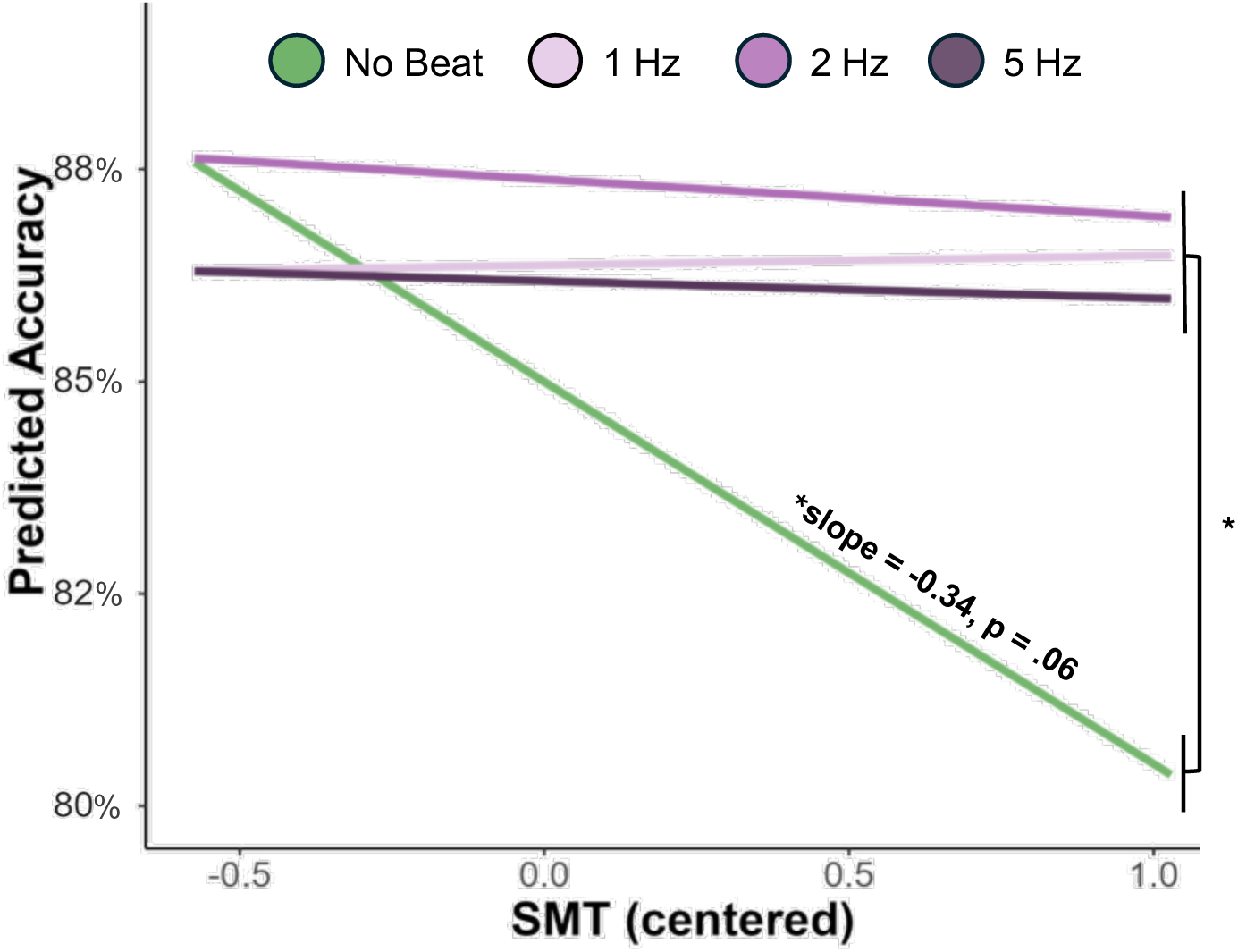
Predicted probability of target detection according to SMT and rhythmic stimulation condition. In the no-beat condition, as the SMT increases (slower participants), the probability of detecting targets decreases. Instead, when there is a beat (1, 2 or 5 Hz), the probability of detecting targets is not dependent on SMT.

In the no-beat condition, we observed a trending monotonic reduction in detection performance as a function of personal tempo (i.e., individuals with slower SMTs generally performed worse; slope estimate = −0.339, p = .065, no-beat condition). In the rhythmic stimulation conditions (1, 2, 5 Hz), we observed no such relationship. Individuals’ detection performance was similar across varying personal tempi (i.e., all slopes ∼0; 1 Hz, slope estimate = 0.0096, p = 0.95; 2 Hz, slope estimate = −0.039, p = 0.83; 5 Hz, slope estimate = −0.017, p = 0.93).

The slopes of those linear trends, in fact, were significantly different between all external rhythms and the no-beat condition. In other words, the presence of an external rhythm significantly impacted the relationship between detection performance and personal tempo (no beat vs. 1 Hz, slope = −0.35, p < .001; no beat vs. 2 Hz, slope = −0.30, p < .005; no beat vs. 5 Hz, slope = −0.32, p < .001; all other comparisons between rhythmic beats, n.s.). While without a beat, individuals with slower spontaneous tempo generally performed worse than those with faster spontaneous tempo, when an external beat was present, detection performance was independent of personal tempo. Essentially, those with a slower spontaneous tempo benefited from rhythmic stimulation (compared to having no beat); while individuals who exhibited a fast spontaneous tempo showed no difference in detection performance between the two conditions (beat vs. no-beat) (see Figure 4, below).

### Performance benefits due to target predictability are unrelated to SMT or rhythmic stimulation

We next assessed whether the target’s predictability affects performance. To this end, we performed a model comparison between the full model and a nested model which excluded the main effect of predictability. We found that the target predictability contributes significantly to the full model, such that detection performance was generally higher for predictable targets, relative to random targets (ΔBIC = 66, χ2 (8) = 22.7, p =.004). Specifically, the odds of detecting predictable targets were 1.09 times higher than the odds of detecting random ones (OR = 1.09, 95% CI: [1.04, 1.14], p <.001; 85.8 ±6.05% vs. 84.8±6.33%). In addition, there were no interactions between predictability and SMT. While a complementary pattern is exhibited in participants’ RTs, target predictability did not contribute significantly to the full model (p =.49, n.s).

## DISCUSSION

The present study examined the impact of external and internal rhythms on detection performance in a non-rhythmic dynamic search task. During the task, participants heard background sounds containing different beats (1-, 2-, 5-Hz, or no beat) which were unrelated to the task and conferred no information about target onset. We found that, overall, auditory rhythmic stimulation enhanced task performance, improving both detection accuracy and response speed. This enhancement was tempo-dependent, with the strongest facilitation observed at 2 Hz. Importantly, the magnitude of this benefit varied with individuals’ spontaneous motor tempo: participants with slower tempi showed performance benefits due to the rhythms, while participants with faster spontaneous tempi performed similarly across all conditions.

In our task, performance benefits occurred regardless of target predictability. Auditory rhythms did not influence the implicit formation of prediction; instead, they exerted a general facilitation on performance. Consistent with a previous study by Boettcher and colleagues (2022), the results show that temporal regularities are incorporated into dynamic priority maps, shaping attention when combined with spatial regularities. Participants were faster and more accurate when both the temporal onset and spatial location of targets were predictable compared to when they were random. However, auditory rhythmic stimulation did not further enhance implicit learning or prediction formation, indicating that the observed facilitation does not arise from improved encoding of temporal structure in the task.

External rhythms within the delta range (0.5–4 Hz) have previously been shown to facilitate attentional processing (Farahbod et al., 2020). This frequency band has been repeatedly linked to temporal attention and top-down anticipatory processes (Haegens & Zion Golumbic, 2018), particularly in auditory processing (Ahissar et al., 2001; Farahbod et al., 2020; Gross et al., 2013; Hickok et al., 2015; Howard & Poeppel, 2010; Lakatos et al., 2013; Large & Jones, 1999; Luo & Poeppel, 2007). However, in the present study, target onsets were not aligned with the background rhythmic stimulation, and target dynamics unfolded over substantially slower timescales (targets faded in and out over 2.5 s). Accordingly, the observed behavioural benefits cannot be explained by precise temporal alignment between the external beat and target occurrence. In our task, auditory rhythmic stimulation may have enhanced tonic alertness or general readiness (Husain et al., 2002; Kiss & Linnell, 2021). This interpretation is consistent with our finding that all rhythmic conditions improved detection accuracy relative to hearing no beat, though it could prove insufficient to fully explain the additional advantage observed for the 2 Hz tempo. Here, we rather propose that the specific benefit of the 2 Hz rhythm may arise because the cognitive system is generally more readily engaged at this timescale. Across domains, this tempo has been identified as a preferred tempo for spontaneous motor behaviour (Hammerschmidt et al., 2021; McAuley et al., 2006; Moelants et al., 2002; Snapiri et al., 2025), auditory perception (Fraisse & Deutsch, 1982; Kliger Amrani & Zion Golumbic, 2020; McAuley & Semple, 1999; Rioux & Grondin, 2025), and sensorimotor synchronisation (McAuley et al., 2006; Repp, 2006; Repp & Su, 2013).

In our task, participants heard auditory rhythmic beats while performing a dynamic search task in the visual modality. It has been shown that auditory stimulation can significantly modulate and enhance visual perception (e.g., Bauer et al., 2021). Early psychophysical studies demonstrated that the presence of a spatially coincident sound can increase the perceived intensity of a visual stimulus (Stein et al., 1996) and facilitate the detection of visual targets (Gleiss & Kayser, 2013; Vroomen & Gelder, 2000). This enhancement can occur even when the sound provides no information about where (McDonald et al., 2000) or when (Gleiss & Kayser, 2014) a target will appear. Studies have found that these cross-modal effects result from involuntary attention to a sound, which enhances perceptual sensitivity (*d’*) to subsequent visual events (Feng et al., 2014; McDonald et al., 2000; McDonald & Ward, 2000).

Individual variability in responsiveness to external rhythms is a well-established phenomenon. Studies in both healthy and clinical populations document considerable heterogeneity in rhythmic facilitation of perception (Bauer et al., 2015; Henry et al., 2025; Lin et al., 2022; Saberi & Hickok, 2021). Similar variability challenges the efficacy of rhythm-based motor rehabilitation therapies (Bella et al., 2017; Leow et al., 2014; Ready et al., 2019). Here we show that this variability is linked to individuals’ spontaneous tempo. Compared to individuals with faster spontaneous tempi, individuals with slower spontaneous tempi (who otherwise performed worse without external temporal structure) especially benefitted from rhythmic stimulation. These findings are consistent with other recent findings showing that rhythmic visual stimulation modulated perceptual discrimination in a similar manner: compared to individuals with faster spontaneous tempi, individuals with slower spontaneous tempi exhibited greater rhythmic facilitation when comparing performance on-versus off-beat (Snapiri et al., 2023).

Given the pervasive presence of rhythms in natural environments, an important future direction is to determine how these effects generalize across sensory modalities. Evidence suggests that the optimal tempo for entrainment differs by modality: periodic temporal attention peaks near 1.4 Hz in audition but closer to 0.7–0.8 Hz in vision (Zalta et al., 2020). Similarly, a recent study observed strongest rhythmic modulation in a visual task following ∼0.77 Hz stimulation (Snapiri et al., 2023).

These findings indicate that interactions between external rhythms and attention are modality-specific. Future studies should examine whether SMT interacts differently with auditory and visual rhythms, and whether in naturalistic multisensory contexts, where multiple rhythmic structures often converge, these rhythms jointly support attentional dynamics.

In conclusion, the present findings demonstrate that external auditory rhythms enhance visual task performance even in the absence of rhythmic structure in the task itself. Rhythms near 2 Hz appear particularly effective, likely because they align both with endogenous delta-range activity and with characteristic motor tempo and perceptual temporal preferences. Together, these results suggest that rhythmic stimulation serves as a powerful scaffold for sustained performance, going beyond structured rhythmic tasks, to more dynamic and temporally unpredictable environments.

## Acknowledgments

We thank Hillel Gottesmann for help with auditory stimuli development and for providing insightful remarks on previous drafts of this manuscript.

## Author Contributions

Conceptualisation – T.S., F.K., A.N.L. Methodology – T.S., F.K., N.S. and A.N.L. Software – T.S., F.K., and N.S. Formal analysis – T.S., F.K., and L.S. Investigation – T.S. Data curation – T.S., F.K., L.S. and N.S. Writing, Original Draft – T.S. and F.K. Writing, Review & Editing – L.S., N.S., and A.N.L. Visualisation – T.S., F.K., and A.N.L. Supervision – F.K. and A.N.L.

## Bibliography

Abikoff, H., Courtney, M. E., Szeibel, P. J., & Koplewicz, H. S. (1996). The effects of auditory stimulation on the arithmetic performance of children with ADHD and nondisabled children. Journal of Learning Disabilities, 29(3), 238–246. 10.1177/002221949602900302

Ahissar, E., Nagarajan, S., Ahissar, M., Protopapas, A., Mahncke, H., & Merzenich, M. M. (2001). Speech comprehension is correlated with temporal response patterns recorded from auditory cortex. Proceedings of the National Academy of Sciences, 98(23), 13367–13372. 10.1073/pnas.201400998

Angwin, A. J., Wilson, W. J., Copland, D. A., Barry, R. J., Myatt, G., & Arnott, W. L. (2018). The impact of auditory white noise on semantic priming. Brain and Language, 180–182, 1–7. 10.1016/j.bandl.2018.04.001

Bauer, A. K. R., Ede, F. Van, Quinn, A. J., & Nobre, A. C. (2021). Rhythmic modulation of visual perception by continuous rhythmic auditory stimulation. Journal of Neuroscience, 41(33), 7065–7075. 10.1523/JNEUROSCI.2980-20.2021

Bauer, A. K. R., Jaeger, M., Thorne, J. D., Bendixen, A., & Debener, S. (2015). The auditory dynamic attending theory revisited: A closer look at the pitch comparison task. Brain Research, 1626, 198–210. 10.1016/j.brainres.2015.04.032

Bégel, V., Seilles, A., & Dalla Bella, S. (2018). Rhythm Workers: A music-based serious game for training rhythm skills. Music and Science, 1. 10.1177/2059204318794369

Bella, S. D., Benoit, C. E., Farrugia, N., Keller, P. E., Obrig, H., Mainka, S., & Kotz, S. A. (2017). Gait improvement via rhythmic stimulation in Parkinson’s disease is linked to rhythmic skills. Scientific Reports, 7. 10.1038/srep42005

Benjamini, Y., & Yekutieli, D. (2001). The control of the false discovery rate in multiple testing under dependency. The Annals of Statistics, 29(4). 10.1214/aos/1013699998

Benoit, C.-E., Dalla Bella, S., Farrugia, N., Obrig, H., Mainka, S., & Kotz, S. A. (2014). Musically cued gait-training improves both perceptual and motor timing in Parkinson’s disease. Frontiers in Human Neuroscience, 8. 10.3389/fnhum.2014.00494

Bharathi, G., Jayaramayya, K., Balasubramanian, V., & Vellingiri, B. (2019). The potential role of rhythmic entrainment and music therapy intervention for individuals with autism spectrum disorders. Journal of Exercise Rehabilitation, 15(2), 180–186. 10.12965/jer.1836578.289

Boettcher, S. E. P., Shalev, N., Wolfe, J. M., & Nobre, A. C. (2022). Right place, right time: Spatiotemporal predictions guide attention in dynamic visual search. Journal of Experimental Psychology: General, 151(2), 348–362. 10.1037/xge0000901

Braun Janzen, T., Koshimori, Y., Richard, N. M., & Thaut, M. H. (2022). Rhythm and Music-Based Interventions in Motor Rehabilitation: Current Evidence and Future Perspectives. Frontiers in Human Neuroscience, 15. 10.3389/fnhum.2021.789467

Cheah, Y., Wong, H. K., Spitzer, M., & Coutinho, E. (2022). Background music and cognitive task performance: a systematic review of task, music, and population impact. Music and Science, 5. 10.1177/20592043221134392

Cochen De Cock, V., Dotov, D., Damm, L., Lacombe, S., Ihalainen, P., Picot, M. C., Galtier, F., Lebrun, C., Giordano, A., Driss, V., Geny, C., Garzo, A., Hernandez, E., Van Dyck, E., Leman, M., Villing, R., Bardy, B. G., & Dalla Bella, S. (2021). BeatWalk: Personalized Music-Based Gait Rehabilitation in Parkinson’s Disease. Frontiers in Psychology, 12. 10.3389/fpsyg.2021.655121

Cravo, A. M., Rohenkohl, G., Wyart, V., & Nobre, A. C. (2013). Temporal expectation enhances contrast sensitivity by phase entrainment of low-frequency oscillations in visual cortex. Journal of Neuroscience, 33(9), 4002–4010. 10.1523/JNEUROSCI.4675-12.2013

Daigmorte, C., Tallet, J., & Astésano, C. (2022). On the foundations of rhythm-based methods in Speech Therapy. Proceedings of the International Conference on Speech Prosody, 2022-May, 47–51. 10.21437/SpeechProsody.2022-10

Desbernats, A., Martin, E., & Tallet, J. (2023). Which factors modulate spontaneous motor tempo? A systematic review of the literature. Frontiers in Psychology, 14. 10.3389/fpsyg.2023.1161052

Engler, B. H., Zamm, A., & Møller, C. (2024). Spontaneous rates exhibit high intra-individual stability across movements involving different biomechanical systems and cognitive demands. Scientific Reports, 14(1), 14876. 10.1038/s41598-024-65788-6

Farahbod, H., Saberi, K., & Hickok, G. (2020). The rhythm of attention: Perceptual modulation via rhythmic entrainment is lowpass and attention mediated. Attention, Perception, and Psychophysics, 82(7), 3558–3570. 10.3758/s13414-020-02095-y

Feng, W., Störmer, V. S., Martinez, A., McDonald, J. J., & Hillyard, S. A. (2014). Sounds Activate Visual Cortex and Improve Visual Discrimination. The Journal of Neuroscience, 34(29), 9817–9824. 10.1523/JNEUROSCI.4869-13.2014

Fraisse, P., & Deutsch, D. (1982). The psychology of music. Presses Universitaires de France.

Fujii, S., & Wan, C. Y. (2014). The role of rhythm in speech and language rehabilitation: The SEP hypothesis. Frontiers in Human Neuroscience, 8(OCT). 10.3389/fnhum.2014.00777

Gleiss, S., & Kayser, C. (2013). Eccentricity dependent auditory enhancement of visual stimulus detection but not discrimination. Frontiers in Integrative Neuroscience, 7. 10.3389/fnint.2013.00052

Gleiss, S., & Kayser, C. (2014). Acoustic Noise Improves Visual Perception and Modulates Occipital Oscillatory States. Journal of Cognitive Neuroscience, 26(4), 699–711. 10.1162/jocn_a_00524

Gross, J., Hoogenboom, N., Thut, G., Schyns, P., Panzeri, S., Belin, P., & Garrod, S. (2013). Speech Rhythms and Multiplexed Oscillatory Sensory Coding in the Human Brain. PLoS Biology, 11(12). 10.1371/journal.pbio.1001752

Haegens, S., & Zion Golumbic, E. (2018). Rhythmic facilitation of sensory processing: A critical review. Neuroscience and Biobehavioral Reviews, 86, 150–165. 10.1016/j.neubiorev.2017.12.002

Hammerschmidt, D., Frieler, K., & Wöllner, C. (2021). Spontaneous Motor Tempo: Investigating Psychological, Chronobiological, and Demographic Factors in a Large-Scale Online Tapping Experiment. Frontiers in Psychology, 12. 10.3389/fpsyg.2021.677201

Henry, M. J., Obleser, J., Crusey, M. R., Fuller, E. R., Lee, Y. S., Meyer, M., Acosta, E. A. M., Van Hedger, S. C., Inbar, M., Oderbolz, C., Dunham, S. A., Anankul, Y., Sabo, L. E., Keitel, C., Maddox, R. K., Mehl, K., Aslan, G., Martens, P. A., Sauppe, S., … Peelle, J. E. (2025). How strong is the rhythm of perception? A registered replication of Hickok et al. (2015). Royal Society Open Science, 12(6). 10.1098/rsos.220497

Herbst, S. K., & Landau, A. N. (2016). Rhythms for cognition: The case of temporal processing. Current Opinion in Behavioral Sciences, 8, 85–93. 10.1016/j.cobeha.2016.01.014

Hickok, G., Farahbod, H., & Saberi, K. (2015). The rhythm of perception: entrainment to acoustic rhythms induces subsequent perceptual oscillation. Psychological Science, 26(7), 1006–1013. 10.1177/0956797615576533

Hine, K., Abe, K., & Nakauchi, S. (2025). Spontaneous motor tempo modulates the effect of music tempo on arousal levels. Psychology of Music, 53(4), 653–665. 10.1177/03057356241311288

Hope, R. M. (2013). Rmisc: Ryan Miscellaneous. *R package version 1.5.1 [computer software]*.

Howard, M. F., & Poeppel, D. (2010). Discrimination of speech stimuli based on neuronal response phase patterns depends on acoustics but not comprehension. Journal of Neurophysiology, 104(5), 2500–2511. 10.1152/jn.00251.2010

Husain, G., Thompson, W. F., & Schellenberg, E. G. (2002). Effects of Musical Tempo and Mode on Arousal, Mood, and Spatial Abilities. Music Perception, 20(2), 151–171. 10.1525/mp.2002.20.2.151

Inbar, M., Grossman, E., & Landau, A. N. (2020). Sequences of Intonation Units form a ∼ 1 Hz rhythm. Scientific Reports, 10(1). 10.1038/s41598-020-72739-4

Inbar, M., Grossman, E., & Landau, A. N. (2025). A universal of speech timing: Intonation units form low-frequency rhythms. Proceedings of the National Academy of Sciences of the United States of America, 122(34). 10.1073/pnas.2425166122

Jaeger, T. F. (2008). Categorical data analysis: Away from ANOVAs (transformation or not) and towards logit mixed models. Journal of Memory and Language, 59(4), 434–446. 10.1016/j.jml.2007.11.007

Jones, M. R. (2010). Attending to sound patterns and the role of entrainment. In Attention and Time (pp. 317–330). Oxford University Press. 10.1093/acprof:oso/9780199563456.003.0023

Kiss, L., & Linnell, K. J. (2021). The effect of preferred background music on task-focus in sustained attention. Psychological Research, 85(6), 2313–2325. 10.1007/s00426-020-01400-6

Kiss, L., Szikora, B., & Linnell, K. J. (2024). Music in the eye of the beholder: a pupillometric study on preferred background music, attentional state, and arousal. Psychological Research, 88(5), 1616–1628. 10.1007/s00426-024-01963-8

Kliger Amrani, A., & Zion Golumbic, E. (2020). Spontaneous and stimulus-driven rhythmic behaviors in ADHD adults and controls. Neuropsychologia, 146, 107544. 10.1016/j.neuropsychologia.2020.107544

Lakatos, P., Gross, J., & Thut, G. (2019). A New Unifying Account of the Roles of Neuronal Entrainment. Current Biology, 29(18), R890–R905. 10.1016/j.cub.2019.07.075

Lakatos, P., Musacchia, G., O’Connel, M. N., Falchier, A. Y., Javitt, D. C., & Schroeder, C. E. (2013). The Spectrotemporal Filter Mechanism of Auditory Selective Attention. Neuron, 77(4), 750–761. 10.1016/j.neuron.2012.11.034

Large, E. W., & Jones, M. R. (1999). The Dynamics of Attending: How People Track Time-Varying Events. Psychological Review, 106(1), 119–159.

Lenth, R., & Piaskowski, J. (2025). emmeans: Estimated Marginal Means, aka Least-Squares Means (R package version 2.0.0).

Leow, L.-A., Parrott, T., & Grahn, J. A. (2014). Individual Differences in Beat Perception Affect Gait Responses to Low- and High-Groove Music. Frontiers in Human Neuroscience, 8. 10.3389/fnhum.2014.00811

Lim, K. B., Kim, Y. K., Lee, H. J., Yoo, J., Hwang, J. Y., Kim, J. A., & Kim, S. K. (2013). The therapeutic effect of neurologic music therapy and speech language therapy in post-stroke aphasic patients. Annals of Rehabilitation Medicine, 37(4), 556–562. 10.5535/arm.2013.37.4.556

Lin, W. M., Oetringer, D. A., Bakker-Marshall, I., Emmerzaal, J., Wilsch, A., ElShafei, H. A., Rassi, E., & Haegens, S. (2022). No behavioural evidence for rhythmic facilitation of perceptual discrimination. European Journal of Neuroscience, 55(11–12), 3352–3364. 10.1111/ejn.15208

Luo, H., & Poeppel, D. (2007). Phase patterns of neuronal responses reliably discriminate speech in human auditory cortex. Neuron, 54(6), 1001–1010. 10.1016/j.neuron.2007.06.004

McAuley, J. D., Jones, M. R., Holub, S., Johnston, H. M., & Miller, N. S. (2006). The time of our lives: Life span development of timing and event tracking. Journal of Experimental Psychology: General, 135(3), 348–367. 10.1037/0096-3445.135.3.348

McAuley, J. D., & Semple, P. (1999). The effect of tempo and musical experience on perceived beat. Australian Journal of Psychology, 51(3), 176–187. 10.1080/00049539908255355

McDonald, J. J., Teder-Sälejärvi, W. A., & Hillyard, S. A. (2000). Involuntary orienting to sound improves visual perception. Nature, 407(6806), 906–908. 10.1038/35038085

McDonald, J. J., & Ward, L. M. (2000). Involuntary Listening Aids Seeing: Evidence From Human Electrophysiology. Psychological Science, 11(2), 167–171. 10.1111/1467-9280.00233

McIntosh, G. C., Brown, S. H., Rice, R. R., & Thaut, M. H. (1997). Rhythmic auditory-motor facilitation of gait patterns in patients with Parkinson’s disease. Journal of Neurology Neurosurgery and Psychiatry, 62(1), 22–26. 10.1136/jnnp.62.1.22

Mendes, C. G., Diniz, L. A., & Marques Miranda, D. (2021). Does Music Listening Affect Attention? A Literature Review. Developmental Neuropsychology, 46(3), 192–212. 10.1080/87565641.2021.1905816

Meteyard, L., & Davies, R. A. I. (2020). Best practice guidance for linear mixed-effects models in psychological science. Journal of Memory and Language, 112. 10.1016/j.jml.2020.104092

Moelants, D., Stevens, C., Burnham, D., McPherson, G., Schubert, E., & Renwick, J. (2002). Proceedings of the 7th international conference on music perception and cognition.

Morey, R. D. (2008). Confidence Intervals from Normalized Data: A correction to Cousineau (2005). Tutorials in Quantitative Methods for Psychology, 4(2), 61–64. 10.20982/tqmp.04.2.p061

Nobre, A. C., & Van Ede, F. (2018). Anticipated moments: Temporal structure in attention. Nature Reviews Neuroscience, 19(1), 34–48. 10.1038/nrn.2017.141

Nombela, C., Hughes, L. E., Owen, A. M., & Grahn, J. A. (2013). Into the groove: Can rhythm influence Parkinson’s disease? Neuroscience & Biobehavioral Reviews, 37(10), 2564–2570. 10.1016/j.neubiorev.2013.08.003

Park, M.-S., Byun, K.-W., Park, Y.-K., Kim, M.-H., Jung, S.-H., & Kim, H. (2013). Effect of complex treatment using visual and auditory stimuli on the symptoms of attention deficit/hyperactivity disorder in children. Journal of Exercise Rehabilitation, 9(2), 316–325. 10.12965/jer.130017

Peirce, J., Gray, J. R., Simpson, S., MacAskill, M., Höchenberger, R., Sogo, H., Kastman, E., & Lindeløv, J. K. (2019). PsychoPy2: Experiments in behavior made easy. Behavior Research Methods, 51(1), 195–203. 10.3758/s13428-018-01193-y

Poeppel, D., & Assaneo, M. F. (2020). Speech rhythms and their neural foundations. Nature Reviews Neuroscience, 21(6), 322–334. 10.1038/s41583-020-0304-4

Puyjarinet, F., Bégel, V., Lopez, R., Dellacherie, D., & Dalla Bella, S. (2017). Children and adults with Attention-Deficit/Hyperactivity Disorder cannot move to the beat. Scientific Reports, 7(1), 11550. 10.1038/s41598-017-11295-w

Ready, E. A., McGarry, L. M., Rinchon, C., Holmes, J. D., & Grahn, J. A. (2019). Beat perception ability and instructions to synchronize influence gait when walking to music-based auditory cues. Gait & Posture, 68, 555–561. 10.1016/j.gaitpost.2018.12.038

Repp, B. H. (2006). Rate Limits of Sensorimotor Synchronization. Advances in Cognitive Psychology, 2(2), 163–181. 10.2478/v10053-008-0053-9

Repp, B. H., & Su, Y.-H. (2013). Sensorimotor synchronization: A review of recent research (2006–2012). Psychonomic Bulletin & Review, 20(3), 403–452. 10.3758/s13423-012-0371-2

Rimoldi, H. J. A. (1951). Personal tempo. The Journal of Abnormal and Social Psychology, 46(3), 283–303. 10.1037/h0057479

Rioux, P. A., & Grondin, S. (2025). A cross sectional investigation of the development of rhythmic preferences with motor and perceptual tests. Scientific Reports, 15(1). 10.1038/s41598-025-87631-2

Roman, I. R., Roman, A. S., Kim, J. C., & Large, E. W. (2023). Hebbian learning with elasticity explains how the spontaneous motor tempo affects music performance synchronization. PLOS Computational Biology, 19(6), e1011154–e1011154. 10.1371/journal.pcbi.1011154

Saberi, K., & Hickok, G. (2021). Forward Entrainment: Evidence, Controversies, Constraints, and Mechanisms. 10.1101/2021.07.06.451373

Schön, D., & Tillmann, B. (2015). Short- and long-term rhythmic interventions: perspectives for language rehabilitation. Annals of the New York Academy of Sciences, 1337(1), 32–39. 10.1111/nyas.12635

Shalev, N., Bauer, A.-K. R., & Nobre, A. C. (2019). The tempos of performance. Current Opinion in Psychology, 29, 254–260. 10.1016/j.copsyc.2019.06.003

Shalev, N., & Nobre, A. C. (2022). Eyes wide open: Regulation of arousal by temporal expectations. Cognition, 224, 105062. 10.1016/j.cognition.2022.105062

Siegelman, N., Bogaerts, L., & Frost, R. (2017). Measuring individual differences in statistical learning: Current pitfalls and possible solutions. Behavior Research Methods, 49(2), 418–432. 10.3758/s13428-016-0719-z

Snapiri, L., Kaplan, Y., & Landau, A. N. (2025). From Individual to Shared Tempo: How Spontaneous Tempo Preferences Impact Joint Performance. 10.1101/2025.09.14.676070

Snapiri, L., Kaplan, Y., Shalev, N., & Landau, A. N. (2023). Rhythmic modulation of visual discrimination is linked to individuals’ spontaneous motor tempo. European Journal of Neuroscience, 57(4), 646–656. 10.1111/ejn.15898

Stein, B. E., London, N., Wilkinson, L. K., & Price, D. D. (1996). Enhancement of Perceived Visual Intensity by Auditory Stimuli: A Psychophysical Analysis. Journal of Cognitive Neuroscience, 8(6), 497–506. 10.1162/jocn.1996.8.6.497

Thaut, M. H., & Abiru, M. (2010). Rhythmic auditory stimulation in rehabilitation of movement disorders: a review of current research. Music Perception, 27(4), 263–269. 10.1525/mp.2010.27.4.263

Thaut, M. H., McIntosh, G. C., & Hoemberg, V. (2015). Neurobiological foundations of neurologic music therapy: rhythmic entrainment and the motor system. Frontiers in Psychology, 5. 10.3389/fpsyg.2014.01185

Thaut, M. H., McIntosh, G. C., & Rice, R. R. (1997). Rhythmic facilitation of gait training in hemiparetic stroke rehabilitation. Journal of the Neurological Sciences, 151(2), 207–212. 10.1016/S0022-510X(97)00146-9

Thaut, M. H., McIntosh, G. C., Rice, R. R., Miller, R. A., Rathbun, J., & Brault, J. M. (1996). Rhythmic auditory stimulation in gait training for Parkinson’s disease patients. Movement Disorders, 11(2), 193–200. 10.1002/mds.870110213

Thompson, W. F., Schellenberg, E. G., & Letnic, A. K. (2012). Fast and loud background music disrupts reading comprehension. Psychology of Music, 40(6), 700–708. 10.1177/0305735611400173

Varlet, M., Marin, L., Issartel, J., Schmidt, R. C., & Bardy, B. G. (2012). Continuity of Visual and Auditory Rhythms Influences Sensorimotor Coordination. PLOS ONE, 7(9), e44082-. 10.1371/journal.pone.0044082

Vroomen, J., & Gelder, B. de. (2000). Sound enhances visual perception: Cross-modal effects of auditory organization on vision. Journal of Experimental Psychology: Human Perception and Performance, 26(5), 1583–1590. 10.1037/0096-1523.26.5.1583

Wiltshire, C. E. E., Cler, G. J., Chiew, M., Freudenberger, J., Chesters, J., Healy, M. P., Hoole, P., & Watkins, K. E. (2024). Speaking to a metronome reduces kinematic variability in typical speakers and people who stutter. PLOS ONE, 19(10), e0309612–e0309612. 10.1371/journal.pone.0309612

Wu, C.-C., & Shih, Y.-N. (2021). The effects of background music on the work attention performance between musicians and non-musicians. International Journal of Occupational Safety and Ergonomics, 27(1), 201–205. 10.1080/10803548.2018.1558854

Yoo, G. E., & Kim, S. J. (2016). Rhythmic ruditory cueing in motor rehabilitation for stroke patients: systematic review and meta-analysis. Journal of Music Therapy, 53(2), 149–177. 10.1093/jmt/thw003

Zalta, A., Petkoski, S., & Morillon, B. (2020). Natural rhythms of periodic temporal attention. Nature Communications, 11(1). 10.1038/s41467-020-14888-8

Zamm, A., Pfordresher, P. Q., & Palmer, C. (2015). Temporal coordination in joint music performance: effects of endogenous rhythms and auditory feedback. Experimental Brain Research, 233(2), 607–615. 10.1007/s00221-014-4140-5

Zamm, A., Wang, Y., & Palmer, C. (2018). Musicians’ natural frequencies of performance display optimal temporal stability. Journal of Biological Rhythms, 33(4), 432–440. 10.1177/0748730418783651

Zamm, A., Wellman, C., & Palmer, C. (2016). Endogenous rhythms influence interpersonal synchrony. Journal of Experimental Psychology: Human Perception and Performance, 42(5), 611–616. 10.1037/xhp0000201

